# Myo-optogenetics: optogenetic stimulation and electrical recording in skeletal muscles

**DOI:** 10.1101/2024.06.21.600113

**Authors:** Jeong Jun Kim, Isis S. Wyche, William Olson, Jiaao Lu, Muhannad S. Bakir, Samuel J. Sober, Daniel H. O’Connor

## Abstract

Complex movements involve highly coordinated control of local muscle elements. Highly controlled perturbations of motor outputs can reveal insights into the neural control of movements. Here we introduce an optogenetic method, compatible with electromyography (EMG) recordings, to perturb muscles in transgenic mice. By expressing channelrhodopsin in muscle fibers, we achieved noninvasive, focal activation of orofacial muscles, enabling detailed examination of the mechanical properties of optogenetically evoked jaw muscle contractions. We demonstrated simultaneous EMG recording and optical stimulation, revealing the electrophysiological characteristics of optogenetically triggered muscle activity. Additionally, we applied optogenetic activation of muscles in physiologically and behaviorally relevant settings, mapping precise muscle actions and perturbing active behaviors. Our findings highlight the potential of muscle optogenetics to precisely manipulate muscle activity, offering a powerful tool for probing neuromuscular control systems and advancing our understanding of motor control.

## Introduction

Precise and flexible movement requires highly-coordinated control of local muscle elements due to the complexity of musculoskeletal anatomies. Observing and disrupting patterns of localized motor output can reveal insights into the neural control of movements. While recent advances in *in vivo* high-density electromyography have provided high-resolution measurements of individual motor units^1,2^, methods for perturbing muscle activity *in vivo* have not kept pace. While electrical stimulation of muscles has been long-established since the bioelectricity studies of Galvani and Volta, it is a method with limited spatial and cell-type specificity. Crucially, it is generally incompatible with electromyography (EMG), precluding simultaneous readout and perturbation of muscle activity.

In contrast, optogenetics offers a noninvasive, EMG-compatible method to induce focal electrical activity in skeletal muscles. Activation of channelrhodopsin (ChR) in the sarcolemma can depolarize myocytes and bypass the physiological patterns of muscle recruitment via motor units^3–5^. This direct activation of muscle fibers therefore uncouples muscle activity from motor neuron activity. Spatial control of the light stimulus permits focal activation of muscle regions. In short, optogenetic perturbation of muscles is a precise and powerful strategy for probing neuromuscular control systems.

Here we demonstrate an optogenetic method compatible with EMG recordings in orofacial muscles of transgenic mice. We investigate the mechanical and electrophysiological properties of channelrhodopsin-activated jaw muscle contractions. Based on these findings, we demonstrate uses of muscle optogenetics in mapping local muscle actions and in perturbing active behaviors in a closed loop fashion.

## Results

Our goals were to 1) determine the properties of jaw movements evoked by direct optogenetic activation of muscles; 2) accomplish simultaneous *in vivo* electromyographic recording and optogenetic stimulation of muscle fibers; 3) demonstrate the utility of optogenetic control of orofacial muscles in physiologically and behaviorally relevant settings.

Emx1^Cre^ driver line has well-known central expression in cortical neurons^6^ and lesser-known peripheral expression in whisker muscles^4^, resulting in optically evoked whisking in Emx1^Cre^; channelrhodopsin transgenic mice. While expression in the whisker pad is known, it is not clear if other orofacial muscles, such as the masticatory muscles of the jaw^7^, may also express channelrhodopsin, thus lending a more general opportunity to optically control orofacial movements. To generate mice expressing channelrhodopsin in skeletal muscles, we crossed mice expressing Cre recombinase under Emx1 regulatory sequence (Emx1^Cre^) with mice with Cre-dependent ChR2(H134R)-EYFP reporter cassette (Rosa26^LSL-ChR2-EYFP^). We then set out to determine the patterns of ChR2 expression in Emx1^Cre/+^; Rosa26^LSL-ChR2-EYFP/+^ mice in the orofacial muscles (Fig. 1a). Lightsheet microscopy of CUBIC-cleared jaw showed ChR2/EYFP expression in masticatory muscles of the jaw – the temporalis, and deep and superficial masseters (Fig. 1b). Notably, ChR2/EYFP expression was not observed in the peripheral nerves. Confocal microscopy in immunolabeled temporalis muscle cross-sections showed ChR2-EYFP expression in the sarcolemma (Fig. 1c,d). A closer examination showed weak fluorescence in honeycomb-like structures similar to previous reports of T tubule expression of ChR2 in muscles^3,5^. In contrast, the digastric muscle, a jaw opening muscle, clearly lacked any ChR2 expression (Fig. 1e, f). Thus, histology showed ChR2 expression in the muscle fibers of masticatory muscles.

**Fig. 1:**
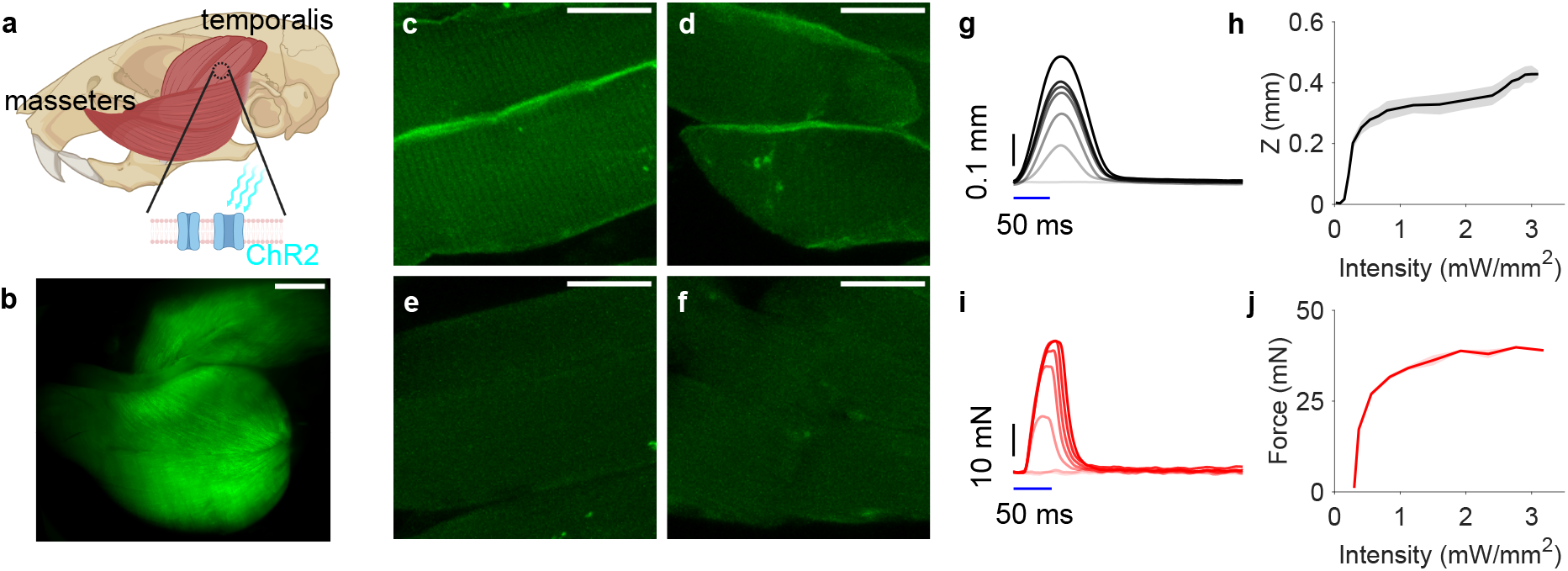
Optogenetic activation of jaw muscles expressing channelrhodopsin. **a** A schematic depicting channelrhodopsin (ChR2) expression in the jaw muscle fibers of Emx1^**Cre/+**^; Rosa26^**LSL-ChR2-EYFP/+**^ mice. **b** Lightsheet images of cleared jaw show ChR2-EYFP (green) expression in temporalis and masseter muscle fibers, absent expression in nerve fibers. Mean projection of 1093 image stack. Scale bar: 2 mm. **c, d** Muscle fibers in temporalis show ChR2-EYFP (green) expression in sarcolemma, as shown in immunolabelled parasagittal (**c**) and coronal (**d**) sections. Scale bar: 20 μm. **e, f** Muscle fibers in digastric muscle do not show ChR2-EYFP expression in parasagittal (**e**) or coronal (**f**) sections. Scale bar: 20 μm. **g** Representative examples of jaw movements induced by 50 ms light pulses of increasing intensity at temporalis. **h** Relationship between light intensity and maximal jaw displacement (Z: jaw closing, n = 5 trials, 1 mouse). Mean ± s.e.m. (error shade). **i** Representative examples of jaw forces under isometric tension induced by 50 ms light pulses of increasing intensity at temporalis. **j** Relationship between light intensity and maximal jaw force (n = 5 trials, 1 mouse). Mean ± s.e.m. (error shade).

We then set out to determine the functional properties of *in vivo* optogenetic control of jaw muscles. Using high-speed (400 fps) videography of the jaw of head-fixed mice, we captured the jaw movements evoked by transcutaneous blue light pulses of varying intensity and duration under anesthesia. Unilateral illumination of the temporalis muscle, a powerful jaw closing muscle, elevated and deviated the mandible from the midline (Supplementary Video 1). The amplitude of the jaw movement in response to 50-ms pulses of increasing light intensity (range 0.02 - 3.1 mW/mm^2^) increased monotonically up to 0.8 mW/mm^2^ (Fig. 1g,h). Similarly, we used a force lever to quantify isometric muscle force production during optically evoked movements under anesthesia. The amplitude of jaw forces in response to 50-ms pulses of increasing light intensity saturated at 39 mN (Fig. 1i, j). In addition, we tested the linearity of forces produced by direct optogenetic activation with a double pulse of optogenetic stimuli separated in varying time intervals. This showed sublinear summation in peak force and impulse at timescales of 25-50ms (Supplementary Fig. 1 f-h), indicating premature contraction at second pulse that interferes with relaxation after first pulse.

To confirm that the jaw movements following optogenetic stimuli are the result of direct activation of muscle fibers, rather than motor axons, we quantified jaw movements during neuromuscular junction block. Curarized jaw muscles showed similar responses to light stimulation, consistent with direct activation of jaw muscle fibers (Supplementary Fig. 1e).

We then recorded the muscle activity in response to optical stimulation. We found the relatively large jaw muscles in mice to be highly amenable to electrophysiological recordings. Here we demonstrate simultaneous *in vivo* EMG recording and optical muscle activation in anesthetized mice using Myomatrix arrays^1^ (Fig. 2, Supplementary Fig. 2).

**Fig. 2:**
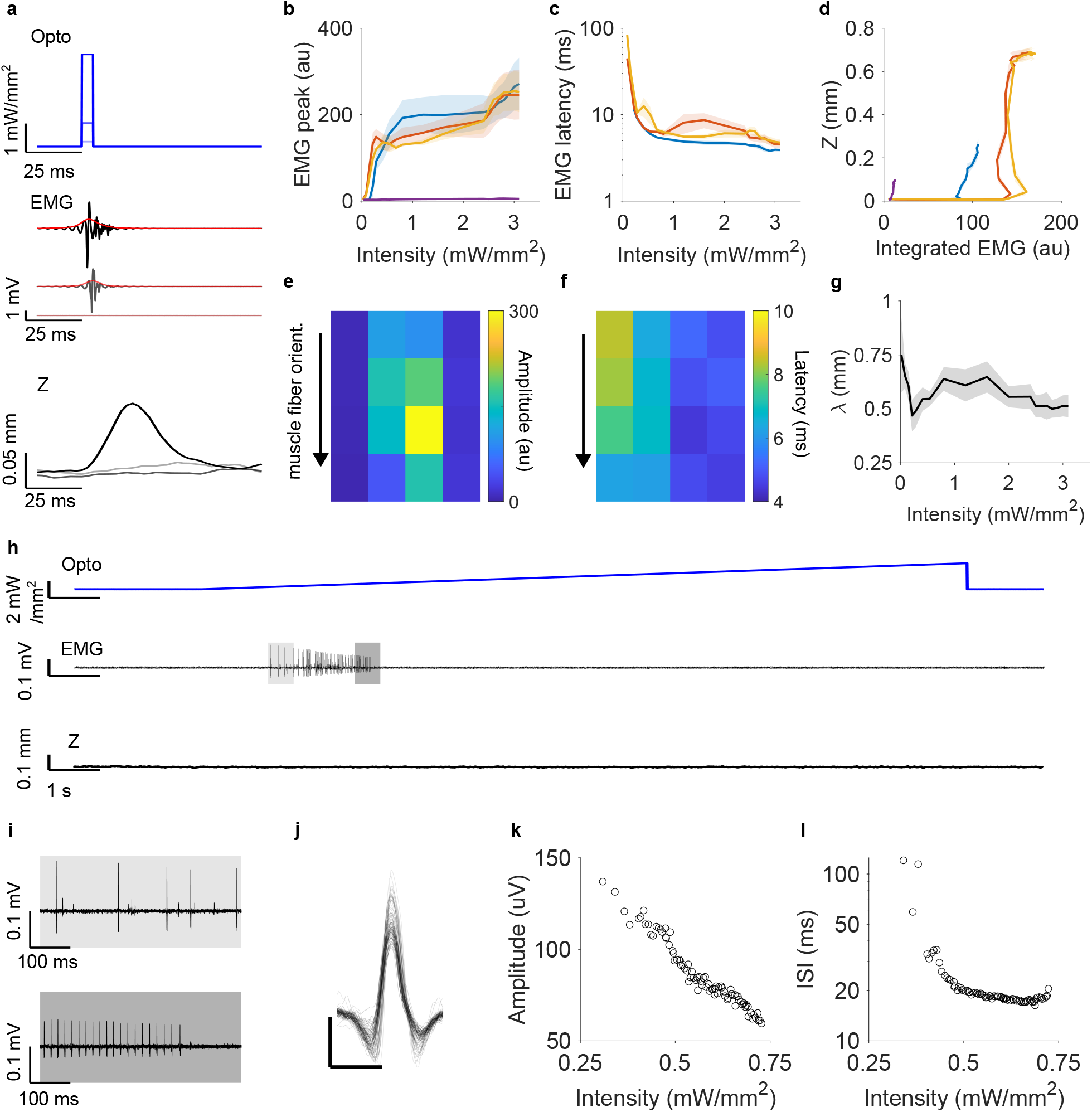
Electromyographic activity in light-activated contractions. **a** Example EMG recording with simultaneous jaw tracking during optogenetic activation of jaw muscles. Top: 5 ms light pulses of varying light intensity; middle: recorded EMG activity (black) and smoothened EMG signal (red); bottom: jaw movements (jaw closing) captured with high-speed videography. **b-d** Relationship between light intensity and peak amplitude of smoothened EMG signal (**b**), between light intensity and latency to peak (**c**), and between sum of smoothened EMG signal and maximum jaw displacement (**d**) in 5 ms (blue), 50 ms (red), 100 ms (yellow) light pulses and 1 Hz sinusoid stimuli (purple) (n = 5 trials per stimulus, 4 recordings, 2 mice). Mean ± s.e.m. (error shade). **e, f** Spatial profile of peak amplitude of smoothened EMG signal (**e**) and latency to peak (**f**) for single trial at 1mW/mm^2^ illumination intensity shows decaying peak amplitude and increasing latency with distance from center of illumination. **g** Space constant of peak amplitude does not show clear dependence on light intensity (n = 5 trials, 1 recording, 1 mouse). **h-l** Properties of EMG signals during slowly ramping optogenetic stimuli. **h** Representative recording shows initial EMG activity followed by suppression without overt jaw movements. Top: light intensity, middle: EMG signal, bottom: jaw movement. **i** Sparse waveforms are observed in the EMG signals during slow ramp (**h**: shaded insets). **j** Spike waveforms colored with increasing opacity based on light intensity shows decreasing amplitude with preserved shape of waveform. **k** Amplitude of spike waveforms decrease with light intensity. **l** Inter-spike interval (ISI) decreases to a plateau at 16 ms with increasing light intensity.

We recorded muscle activity induced by optogenetic stimuli under varying parameters. In pulsed stimuli, increasing the light intensity increased the amplitude of the EMG signal, reflecting increased activation of muscle fibers and greater jaw displacement (Fig. 2a, b, d). In addition, increasing the light intensity reduced the latency of peak EMG response, reflecting faster depolarization (Fig. 2c). In wild-type mice lacking channelrhodopsin expression in muscles, optogenetic stimulation did not induce electrical activity (Supplementary Fig. 2d), confirming that optically induced activity does not represent light-induced artifacts.

Leveraging the defined layout of the Myomatrix electrode contact sites (Supplementary Fig. 2a, b), we measured the spatial profile of light-evoked electrical activity. Peak EMG activity decayed with distance with a space constant ranging from 0.5 to 0.7 mm (Fig. 2g). The spatial profile of peak EMG activity was anisotropic and oriented along the direction of the temporalis muscle fibers (Fig. 2e). Similarly, the spatial profile of latency to EMG peak showed increasing latency with distance from the center of illumination (Fig. 2f).

Paradoxically, slowly ramping stimuli suppressed muscle activity. We delivered 15 second long ramp illumination stimuli with peak light intensity sufficient to cause large jaw movement in pulsed delivery. This failed to evoke any observable jaw movements (Fig. 2h). During the ramp stimulus, sparse EMG signals were present at early, low light intensity (Fig. 2i, j). The amplitude of the waveforms decreased threefold as ramped light intensity increased (Fig. 2k). The inter spike interval (ISI) of the waveforms did not change greatly after reaching a set ISI of 16.33 ms (Fig. 2l). Beyond the threshold of 0.73 mW/mm^2^, high light intensity suppressed EMG signals. The paradoxical suppression of muscle activity in ramps was consistent with a supercritical Andronov-Hopf bifurcation^8^ in a Hodgkin-Huxley model of muscle action potentials (Supplementary Fig. 3d-h, Supplementary Text).

Finally, we sought to apply optogenetic control of muscle movements in physiologically and behaviorally relevant settings. Muscle optogenetics can be used to map physiologic muscle actions in a noninvasive manner. In anesthetized animals, we delivered optogenetic stimulation to various areas of the jaw musculature, such as the temporalis, superficial masseter, and deep masseter, and recorded resulting movement responses.

Light (50 ms pulses, 1 mW/mm^2^) delivered over both the temporalis and masseter muscles drove elevation of the mandible (Fig. 3a, top). However, optogenetic stimulation of the temporalis deviated the mandible ipsilaterally while optogenetic stimulation of masseter muscles deviated contralaterally. This was evidenced by a rapid transition between illumination sites causing ipsilateral versus contralateral deviation corresponding to the boundaries of the muscles (Fig. 3a, middle). Optogenetic stimulation of the temporalis did not protract or retract the mandible, while optogenetic stimulation of the masseter induced protraction (Fig. 3a, bottom).

**Fig. 3:**
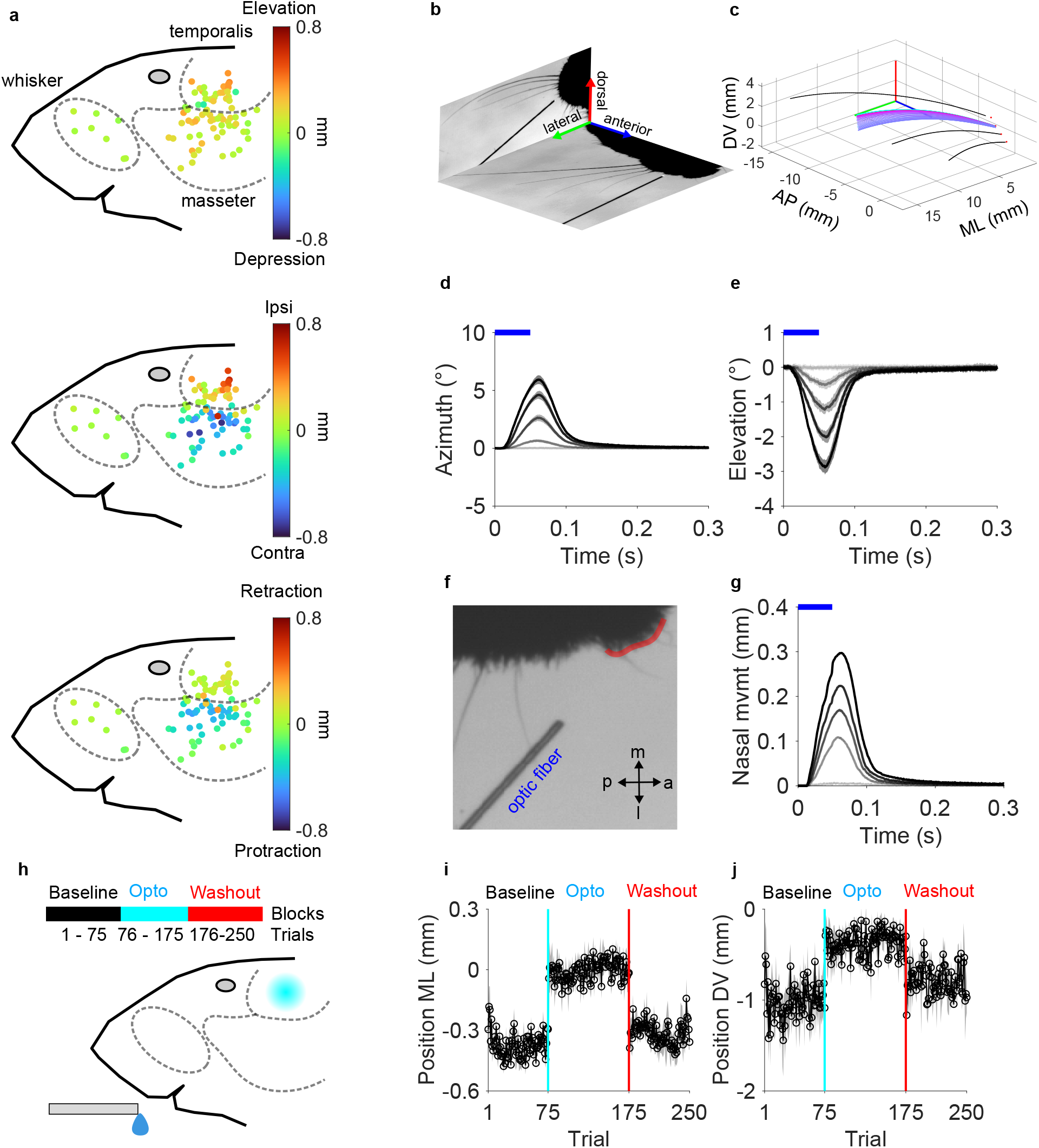
Applications of muscle optogenetics in orofacial areas. **a** Localized activation of jaw muscles with light pulses (1 mW/mm^2^, 50 ms) shows the different actions by temporalis and masseter muscles on the mandible. Temporalis elevates and deviates the mandible ipsilaterally while masseter elevates, protracts, and deviates the mandible contralaterally (n = 4 trials, 2 sessions, 2 mice). **b-e** Optogenetic activation of whisker muscles. **b** High speed videography at 4kHz of whiskers with two camera views. **c** Tracking and 3D reconstruction of C row whiskers with C2 whisker highlighted (color shade: history trace in 1.25 ms intervals). **d, e** Mean traces of C2 whisker protraction **d** and downward deflection **e** induced by light pulses (50 ms, 0 - 3.2 mW/mm^2^, n = 25 trials, 1 mouse). Mean ± s.d. (error shade) **f, g** Optogenetic activation of rhinarial muscles. **f** High-speed videography at 4kHz of nostrils (arrow bar: 5 mm). **g** Mean traces of nasal movement induced by light pulses (50 ms, 0 - 3.2 mW/mm^2,^ n = 5 trials, 1 mouse). Mean ± s.d. (error shade). **h-j** Closed loop optogenetic perturbation of jaw muscles during head-fixed licking behavior. **h** A schematic depicting optogenetic activation of left temporalis muscle (3.2mW/mm^2^, 100 ms pulses) during licks to a water-dispensing lick port. Trial blocks consist of baseline, optogenetic perturbation (opto), and washout blocks. **i-j** Average jaw position in medial-lateral axis (+: ipsilateral deviation) and dorsal-ventral axis (+: elevation) during licks in the baseline, perturbation, and washout blocks does not show motor adaptation (n = 2 mice, 5 sessions). Mean ± s.e.m. (error bar)

In addition, we extended optogenetic activation to other orofacial muscles, including whisker and nasal muscles. Focal, noninvasive optogenetic stimulation is particularly useful for activating small muscles for fine movements. Using high-speed videography at 4000 fps, we captured the whisker and nasal movements evoked by light pulses (50 ms pulses, 0 - 3.2 mW/mm^2^) in anesthetized mice (Fig. 3b, c, f; Supplementary Video 2, 3). Illumination of the rostral whisker pad evoked whisker protractions in anesthetized mice (Fig. 3d), consistent with a previous report^4^. Three-dimensional tracking of whiskers also revealed downward deflections of whiskers (Fig. 3e). Amplitude of whisker movements increased with light intensity. Illumination of the dilator nasi muscle evoked nose movements that increased with light intensity (Fig. 3g). Thus, muscle optogenetics evoked facial movements in the jaw, whiskers, and nostrils.

Finally, we asked if optogenetic muscle stimulation could perturb ongoing movement in active animals. We optogenetically activated jaw muscles (50 ms pulses, 3.2 mW/mm^2^) during licking. Head-fixed mice were trained to lick a water port during behavior trials. Each trial consisted of an auditory cue followed by water reward given after five lick touches to the lick port. After a baseline block of trials, mice proceeded through a ‘perturbation’ block in which the temporalis muscle was optogenetically stimulated during the approximate opening phase of the fourth lick with the stimulation triggered in a closed-loop manner by the port contact of the previous lick (Fig. 3h). The optogenetic perturbation elevated and deviated the jaw from the midline (Fig. 3i, j). Jaw kinematics during the perturbation block did not show motor adaptation between early and late trials. Trials following the perturbation block (“wash-out” block) did not show any lasting motor adaptation. Motor perturbations with direct optogenetic muscle activation were stable across trials without apparent motor adaptation.

## Discussion

In this study, we demonstrate a noninvasive, optical method to focally stimulate or inhibit orofacial muscles with high spatiotemporal resolution compatible with electrophysiological recordings. We characterized optogenetically evoked jaw movements, recorded *in vivo* muscle activity in response to optogenetic stimulation, and applied muscle optogenetics in physiologically and behaviorally relevant settings.

Transcutaneous optogenetic stimulation at physiologically nonharmful light intensity drives robust jaw movements in passive and actively moving animals. Focal, unilateral muscle activation was sufficient to drive movements of behaviorally relevant magnitude (up to ∼0.5 mm, whereas awake licking involves jaw movement of ∼2-3 mm). The optogenetic activation of muscles likely involves asynchronous recruitment of muscle fibers. The asynchronicity contrasts with the orderly recruitment of well-defined groups of muscle fibers or motor units occurring during physiological contractions, as evidenced by the drastic differences in waveforms in optically evoked movements and in spontaneous activity (Supplementary Fig. 2c). The optogenetic activation of muscle fibers likely occurs in a depth-dependent manner, given the highly light absorbing and scattering properties of muscle. The measured spatial constant likely reflects a combination of light scattering and spatial propagation of depolarization. Well-isolated waveforms observed in ramping light stimuli at low intensities likely reflect the sparse, synchronized firing of the most superficial muscle fibers.

Properties of EMG signals were consistent with a simple two-compartment Hodgkin-Huxley model of muscle action potentials (Supplementary Fig. 3, Supplementary Text). In particular, the paradoxical suppression of electrical activity in ramp stimuli was consistent with a nonlinear oscillator undergoing a bifurcation from a stable limit cycle to a stable point of equilibrium^8^.

Paradoxical suppression in slow ramps shows that optogenetic activation or inhibition is dependent on the interaction between channelrhodopsin conductance and electrophysiological properties of muscle fibers. Activating channelrhodopsin in muscles can paradoxically inhibit muscle contractions. The mechanism of suppression is akin to depolarizing neuromuscular blockers, in which persistent depolarization prevents muscle contractions^9^. This observation suggests clinical applications in treating abnormal muscle tone, such as muscle spasticity or atrophy. A single opsin could be used to modulate muscle tone bidirectionally with different stimulus parameters.

In our study, we applied muscle optogenetics to orofacial muscles controlling the whiskers, nose, and jaw. We recorded both the electrical activity and movements generated by optogenetic activation of muscles. As advances in electromyographic technologies provide new methods to “read” muscle activity, optogenetics provide a method to “write” muscle activity simultaneously. Focal activation of nearby sites can cause distinct movements, highlighting the potential of this strategy to drive precise, complex kinematic manipulations. Simultaneous readout and manipulation of muscle activity can be a powerful combination to investigate neural control of movements, implement closed-loop control of motor actions, and study proprioceptive feedback during active movements. We expect that developments of integrated tools to “read” and “write” muscle activity will advance our understanding of motor control.

## Methods

### Subjects

We crossed Emx1-IRES-Cre knock-in homozygotes (Stock no. 005628, Jackson Laboratory) with Rosa26^LSL-ChR2-EYFP/+^ (Ai32) reporter homozygotes (Stock no. 024109, Jackson Laboratory) to obtain Emx1^Cre/+^; Rosa26^LSL-ChR2-EYFP/+^. Emx1, a homeobox gene, is known to express in embryonic pharyngeal arches (http://www.informatics.jax.org/), which give rise to head and neck structures.

### Histology

We transcardially perfused Emx1^Cre/+^; Rosa26^LSL-ChR2-EYFP/+^ mice with PBS followed by 4% PFA in 0.1 M PB, and either dissected jaw musculature for confocal imaging or preserved whole intact tissues for clearing. The tissue was fixed in 4% PFA at least overnight. For confocal imaging of sections, dissected jaw muscles were stored in 30% sucrose in PBS at 4°C, embedded in OCT, flash-frozen, and then sectioned at 20 um thickness with a Leica Cryostat CM3050S. The sections were blocked with 5% bovine serum albumin (BSA) in PBST (0.1% Tween in PBS) for 1 hr. Incubation with primary antibody chicken anti-GFP IgY (Thermo Fisher A10262, 1:1000) lasted overnight, whereas incubation with secondary antibody goat anti-chicken IgG conjugated to Alexa Fluor 488 (Thermo Fisher A-11309, 1:500) lasted 2 h. The sections were mounted with Aqua-Poly/Mount (Polysciences). Images were acquired with a Zeiss LSM 700 confocal scanning microscope.

### Whole tissue clearing and lightsheet imaging

Intact heads were processed for whole tissue clearing with CUBIC protocol. Following previously described CUBIC protocol^10^, samples were delipidated in CUBIC-L solution at 37°C for 4-7 days, washed overnight in PBS, and refractive index matched in CUBIC-R at room temperature for 2-3 days. Samples were oriented such that the imaging plane aligned with the parasagittal planes. Images were acquired with a LaVision BioTec UltraMicroscope II with a large field of view 11.1mm X 13.2 mm and resolution of 5.12 μm/pixel at 3 μm Z steps.

### Optogenetic stimulation and jaw tracking in anesthetized mice

All procedures described below were approved by The Institutional Animal Care and Use Committee at Johns Hopkins University. Mice were implanted with a skull headbar and ground pin over the right visual cortex. The left cheek overlying jaw muscles were depilated with hair removal lotion.

In jaw kinematics experiments, high-speed videography recorded orofacial movements in head-fixed animals under isoflurane anesthesia (0.9-1.5% at flow rate 1L/min). A high-speed CMOS camera (PhotonFocus DR1-D1312-200-G2-8) in a 0.25X telecentric lens (Edmund Optics) captured jaw kinematics at 400 frames per second using IR illumination at 850nm and an angled mirror to collect side and bottom views simultaneously. We tracked eleven keypoints along the jaw using DeepLabCut^11^ and quantified jaw movements.

Similarly, in jaw force experiments, a high precision force lever recorded the forces caused by jaw muscle contraction in head fixed animals under ketamine/xylazine anesthesia (IP injection, ketamine 87.5 mg/kg, xylazine 12.5 mg/kg). A nylon thread yoked the lower incisors to a muscle lever (Aurora Scientific 300C-LR) operating in isometric (length controller) mode. Force signals were digitized and recorded via National Instruments DAQ system (BNC-2090A) and WaveSurfer (HHMI Janelia Research Campus).

Optogenetic stimuli were provided by a 473 nm DPSS laser (MBL-III-473-100, Ultralasers) and delivered to the cheek via a Ø200 um 0.39 NA optical fiber manually positioned around 1 mm away from the surface with intensity ranging from 0-3.2 mW/mm^2^. Optogenetic stimuli consisted of either 1) 1Hz pulse trains of varying pulse width (5, 50, 100 ms) and light intensity, 2) a 1Hz sinusoid with increasing light intensity, or 3) a slow ramp of 15 s that reached the max intensity of 3.2 mW/mm^2^. Stimulus parameters were controlled by a National Instruments DAQ system (BNC-2090A) with WaveSurfer.

For curare experiments, we injected 10mM (+)-tubocurarine hydrochloride (Sigma-Aldrich) in normal saline in 50 μL volumes subcutaneously at the temple. Saline injections served as the control. The curare dosage effective for localized muscle paralysis was confirmed with injections in whiskerpad and visual inspection for asymmetric whisker movements.

### EMG recordings

In a subset of mice, we simultaneously recorded epimysial EMG activity in temporalis muscles with high-speed video and optogenetic stimulation. As previously demonstrated^1^, we implanted Myomatrix electrode arrays (4 × 8 grid) on the temporalis muscle using the epimysial method. The transparent polyimide substrate allowed light delivery through the electrode array. The implanted ground pin served as the ground electrode. We recorded EMG signals in 16 channel bipolar headstage (Intan RHD2216) and band-passed the signals between 300-9000 Hz. The EMG signals were rectified and smoothed with a Gaussian kernel with standard deviation of 2.5ms. Space constant of peak EMG activity was determined with a single exponential fit in MATLAB (fit toolbox).

### Optogenetic stimulation and whisker tracking in anesthetized mice

Recordings of photostimulation-induced whisker and nose motion were performed in two head-fixed mice under ketamine/xylazine anesthesia (IP injection, ketamine 87.5 mg/kg, xylazine 12.5 mg/kg). Prior to recording, the left-side facial fur and all whiskers other than C1 through C4 were trimmed close to the skin to minimize occlusion of the spared whiskers.

High-speed video was recorded under IR illumination (whiskers backlit by an 850nm LED) using an Optronis Cyclone-2-2000 CoaxPress camera, acquiring 4,000 frames per second over a 1216 × 600 pixel field of view with an exposure time of 90 microseconds. Top-down and rear views were acquired simultaneously in a single field of view, using mirrors positioned near the mouse face. The spatial resolution of the video was 25 pixels/mm.

Photostimulation was delivered in 0 - 7.5 mW pulses of 5, 50, and 100ms duration from a DHOM-L-473-200 473nm laser, through an optic fiber positioned within millimeters of the mouse’s face. Photostimulation was targeted to three locations: to the follicle of whisker D5 and dorsal and caudal to whisker C1 (to induce whisker deflection, in one mouse), and rostral to whisker C5 (to induce nose motion, in one mouse).

The four spared whiskers were traced automatically in each view using the Janelia whisker tracker^12^. Whisker points in the images were triangulated to 3D using the direct linear transform (DLT) algorithm and a metric stereo calibration of the imaging system^13^, after using the calibrated epipolar geometry to identify corresponding pairs of points on the 2D whisker traces between the two views. Whisker curves were fitted to the triangulated points as degree 5 polynomials, in an extension to 3D of the analytical approach presented by Pammer et al.^14^, and the whisker base azimuth and elevation angles were derived from these fitted curves. The contour of the whisker pad (if the whiskers were tracked) or nose (if the nose was tracked) in each view was obtained from a spline curve fitted to the silhouette of the mouse face in that view, as previously described^15^. To reduce the effect of tracking error near the face on the whisker base angle, the whisker pad contours in each view were transformed based on the calibrated epipolar geometry of the imaging system into isosurfaces analogous to the 2D face masks previously described^14,15^. The whisker base was estimated as the point 1 mm in the direction of the whisker tangent beyond the intersection of the whisker with the union of the two face mask isosurfaces, and the azimuth and elevation angles of the whisker tangent at that intersection point were taken as the whisker base angles.

### Behavioral task with closed loop optogenetics

Mice were restricted to 1 mL/day of water for ≥ 3 days prior to training. Following a 0.1 s long auditory cue of 15 kHz pure tone, head-fixed mice licked for water at a lick port with water droplets of 2uL given after five lick contacts, detected by a conductive lick detector (Svoboda Lab, HHMI Janelia Research Campus). An Arduino-based system (Teensy 3.2 and Teensyduino) and custom MATLAB-based software served as the task controller. In a behavioral session, trials were separated into 3 blocks of 75 baseline trials without optogenetic stimuli, 100 perturbation trials with closed loop optogenetic stimuli, and 75 washout trials without optogenetic stimuli. For closed loop optogenetics, light pulses of 100 ms at 3.2mW/mm^2^ were triggered 75 ms after the detection of the third lick contact. Following the behavioral session, mice were placed under isoflurane anesthesia, and identical optogenetic stimuli were delivered in the passive setting to confirm optogenetic activation of the muscle. Keypoints along the jaw were tracked with DeepLabCut^11^.

### Biophysical model of muscle action potentials

We constructed a two-compartment Hodgkin-Huxley model for muscle action potentials, corresponding to the sarcolemma and T tubule, with the following pair of equations^16^.

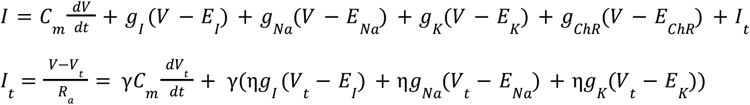

where V, C_m_ are the membrane voltage and capacitance of the sarcolemma, g_L_, g_Na_, g_K_, g_ChR_ the leak, sodium, potassium, and channelrhodopsin conductances, and E_L_, E_Na_, E_K_, and E_ChR_ the respective reversal potentials. I_t_, V_t_ are the T tubule current and membrane voltage, and R_a_ is the access resistance between extracellular space and T tubule. γ denotes the ratio of surface ratio between T tubule and sarcolemma, and η is the relative density of respective ion channels in T tubules compared to sarcolemma. The voltage terms V and V_t_ follow the voltage-dependence in standard Hodgkin-Huxley equations with m and h as the sodium gating variables and n as the potassium gating variable^17^.

Relevant electrophysiological parameters were adapted from previous work (pulses simulation, Cannon et al^16^; sinusoid and ramp simulation, Metzger et al^18^). As ChR2 is a nonselective cation channel, channelrhodopsin reversal potential E_ChR_ was set to 0 mV, and only channelrhodopsin expression in the sarcolemma was considered. The range of channelrhodopsin conductance g_ChR_ was empirically set to 0.05 - 0.5 mS/cm^2^ with maximum value in the same order of magnitude as leak current conductances. The differential equations were numerically integrated with Euler’s method with a time step of 1 μs.

## Supplemental Text

### Hodgkin-Huxley model of action potentials in muscles

With these observations of optical evoked muscle activity, we set out to explain the observed properties with a simple biophysical model. To this end, we created a two-compartment Hodgkin Huxley model of muscle action potentials^16–19^. The model incorporated voltage-dependent and independent currents and optogenetic currents with two electrical compartments corresponding to the T-tubules and sarcolemma (Methods).

The Hodgkin-Huxley model of action potentials recapitulated important observations from EMG recordings. Simulated action potentials showed after depolarizations which are consistent with previous studies of action potentials in ex vivo muscles. In simulated action potentials, the peak membrane voltage increased with light intensity and the latency decreased with light intensity. The effect of the after depolarization was clear in cases of pulses of varying width in which increasing the pulse width did not induce additional action potentials in a single muscle fiber. Moreover, the decrease in latency of the action potentials was evident in the simulations of sinusoidal optogenetic stimuli in which increasing the light intensity led to an earlier onset of action potentials.

To explain the paradoxical suppression of muscle electrical activity in ramping stimuli, we simulated slowly ramping optogenetic stimuli in the Hodgkin-Huxley model.

The simulated action potential showed qualitatively similar behavior as the EMG recordings, in which strong optogenetic currents suppressed action potentials. Trains of action potentials were present at low light intensities, and then terminated upon reaching a critical point in light intensity. At the critical point, as shown in Supplementary Fig. 3e, the action potentials were dampened into arbitrarily small oscillations. The slow ramp simulation exhibited supercritical Andronov-Hopf bifurcation, characterized by the disappearance of a stable limit cycle into a stable equilibrium^8^. The supercritical Andronov-Hopf bifurcation characteristically shows oscillations with vanishing amplitudes, consistent with decreasing waveform amplitudes in EMG recordings, without change in oscillation frequency, consistent with stable inter spike intervals. The disappearance of a stable limit cycle in supercritical Andronov-Hopf bifurcation was reflected in the gating parameters m, h, and n, in which oscillations of decreasing amplitude converged into an equilibrium.

Thus, the Hodgkin Huxley model of muscle action potential recapitulated many of the observations in EMG recordings, particularly the paradoxical suppression of muscle activity in slowly ramping optical stimuli.

## Acknowledgements

JJK was supported by NIH Award 1F31DE033256. WO was supported by Kavli Neuroscience Discovery Institute. SJS was funded by National Institutes of Health Grant R01 NS109237 and U24 NS126936, the McKnight Foundation, and the Simons Foundation as part of the Simons-Emory International Consortium on Motor Control.

---

**Supplementary Fig. 1:**
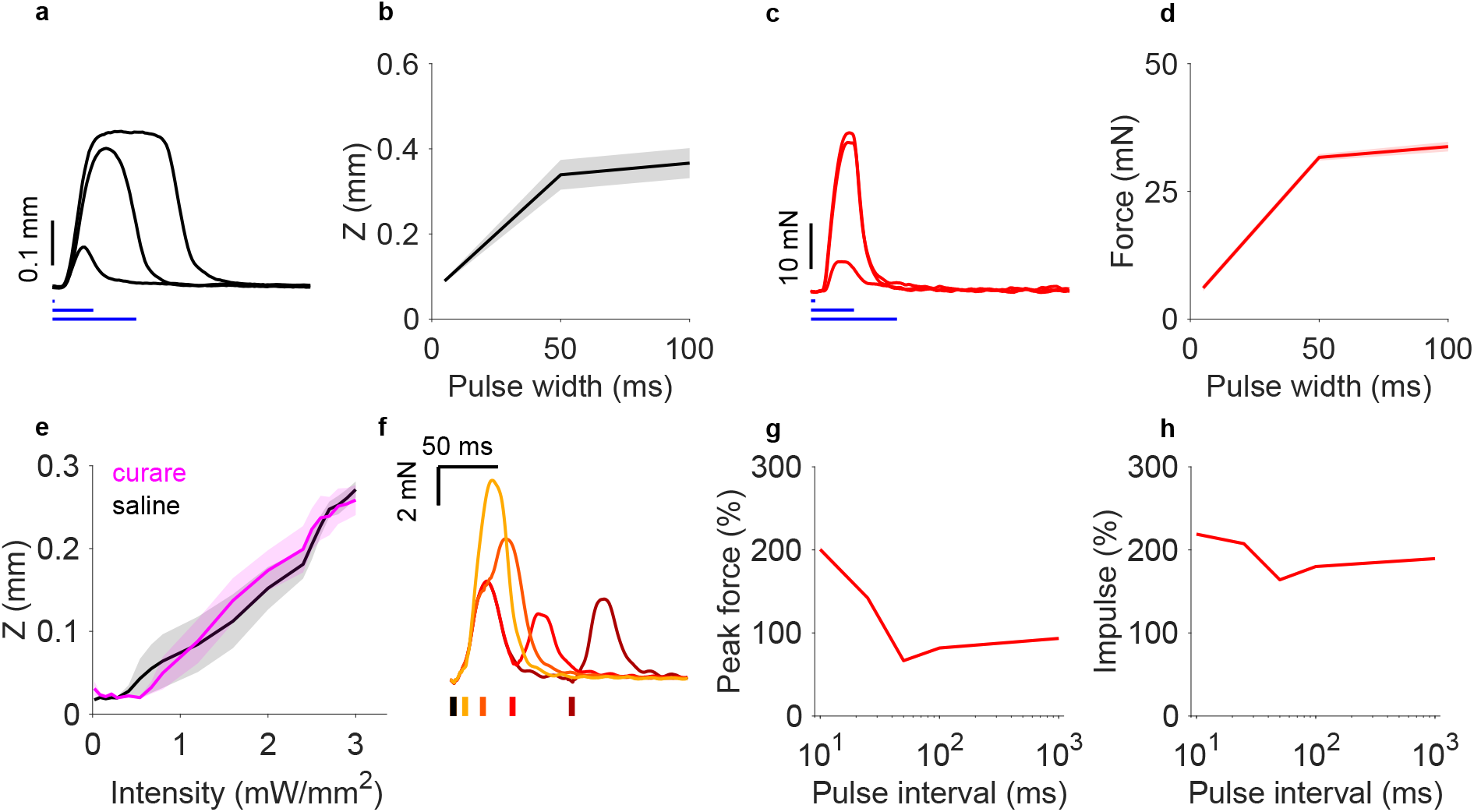
Properties of optogenetically evoked jaw movements. **a** Representative examples of jaw movements induced by light pulses (1mW/mm^**2**^) of varying pulse duration (5, 50, 100 ms) at temporalis. **b** Relationship between pulse duration and maximal jaw displacement (Z: jaw closing, n = 5 trials, 1 mouse). Mean ± s.e.m. (error shade). **c** Representative examples of jaw forces under isometric tension induced by light pulses (1mW/mm^**2**^) of varying pulse duration (5, 50, 100 ms) at temporalis. **d** Relationship between pulse duration and maximal jaw force (n = 5 trials, 1 mouse). Mean ± s.e.m. (error shade). **e** Comparison of curare and saline injection at temporalis for jaw movements under induced by 50 ms light pulses of increasing intensity (n = 5 trials, 2 mice). Mean ± s.e.m. (error shade). Curare does not affect optogenetically evoked muscle movements. **f-h** Linearity of jaw forces in two consecutive pulses of optogenetic activation. **f** Representative examples of jaw forces under isometric tension induced by pairs of light pulses (1mW/mm^**2**^, 5 ms) separated by varying intervals (10, 25, 50, 100 ms) indicated by color shading. **g** Relationship between pulse interval and normalized peak force, defined as the ratio between peak force amplitude from second pulse and peak force amplitude from a single, isolated pulse (n = 5 trials, 1 mouse). Mean ± s.e.m. (error shade). Sublinear summation of muscle forces is present at 25-50 ms pulse interval. **h** Relationship between pulse interval and normalized impulse, defined as the ratio between integral of force produced by two pulses over time and integral of force produced by a single, isolated pulse (n = 5 trials, 1 mouse). Mean ± s.e.m. (error shade).

**Supplementary Fig. 2:**
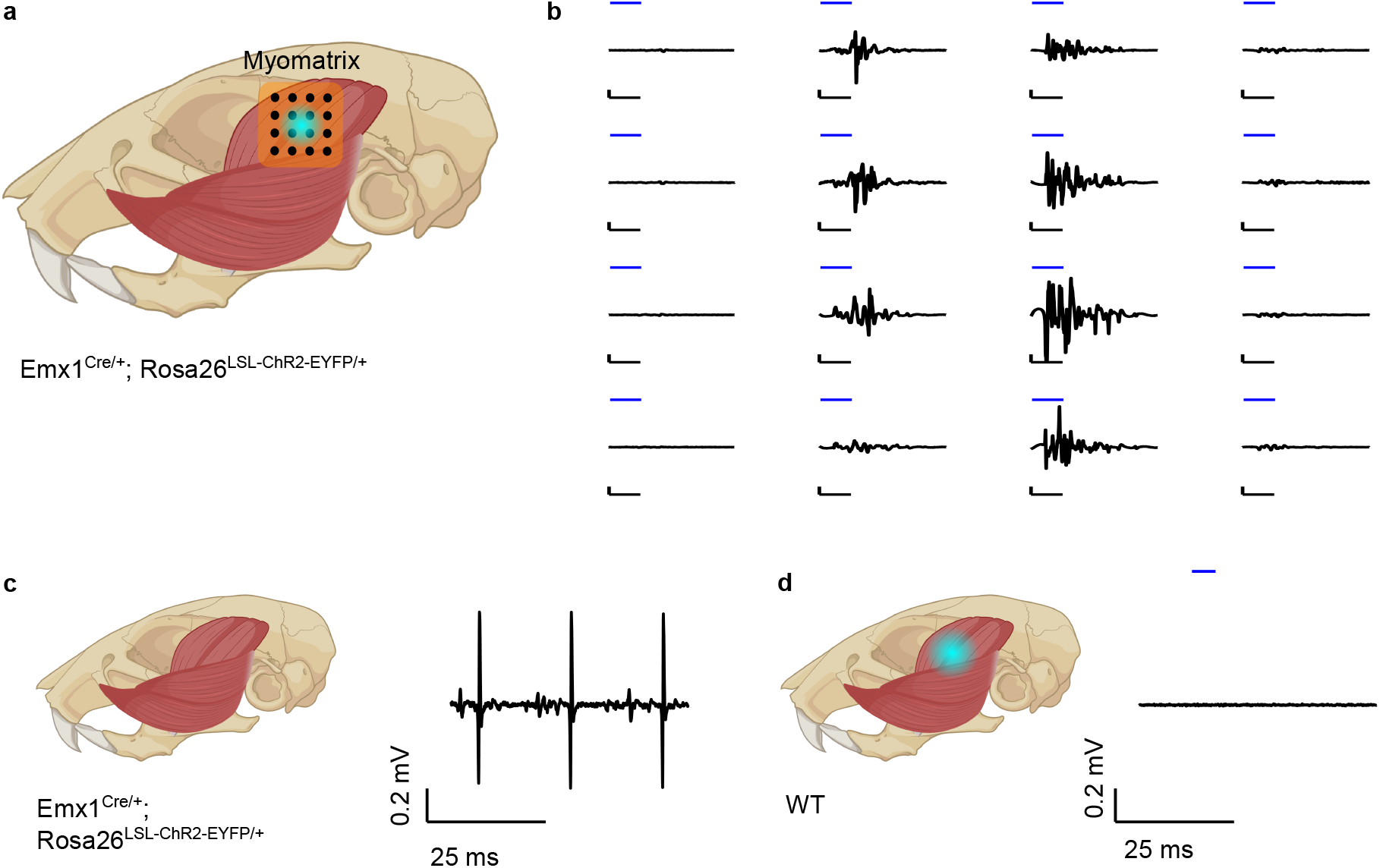
Example electromyography activity during optogenetically evoked and spontaneous movements. **a** Schematic depicting Myomatrix EMG recording in temporalis with optogenetic stimuli. A grid of electrode contacts in a Myomatrix array allows 16 bipolar recording sites. **b** Example EMG signals from temporalis muscle at 16 recording sites during light pulse (1mW/mm^2^, 5 ms). Amplitude of EMG activity decayswith distance from the center of optogenetic stimulation. Vertical scale bar: 0.2 mV; horizontal scale bar: 5 ms. **c** Example spotaneous EMG activity (right) in temporalis muscle of Emx1^Cre/+^; Rosa26LSL-ChR2-EYFP/+ mice (left). Motor unit activity is characterized by sparse, well-defined waveforms in contrast to dense activity during optogenetic stimulation. **d** Example EMG recording (right) in temporalis muscle of wild-type (C57BL/6) mouse during light pulse (3.2 mW/mm^2^, 5ms) (left). Muscle fibers without ChR2 do not show electrical activity in response to optogenetic stimulation.

**Supplementary Fig. 3:**
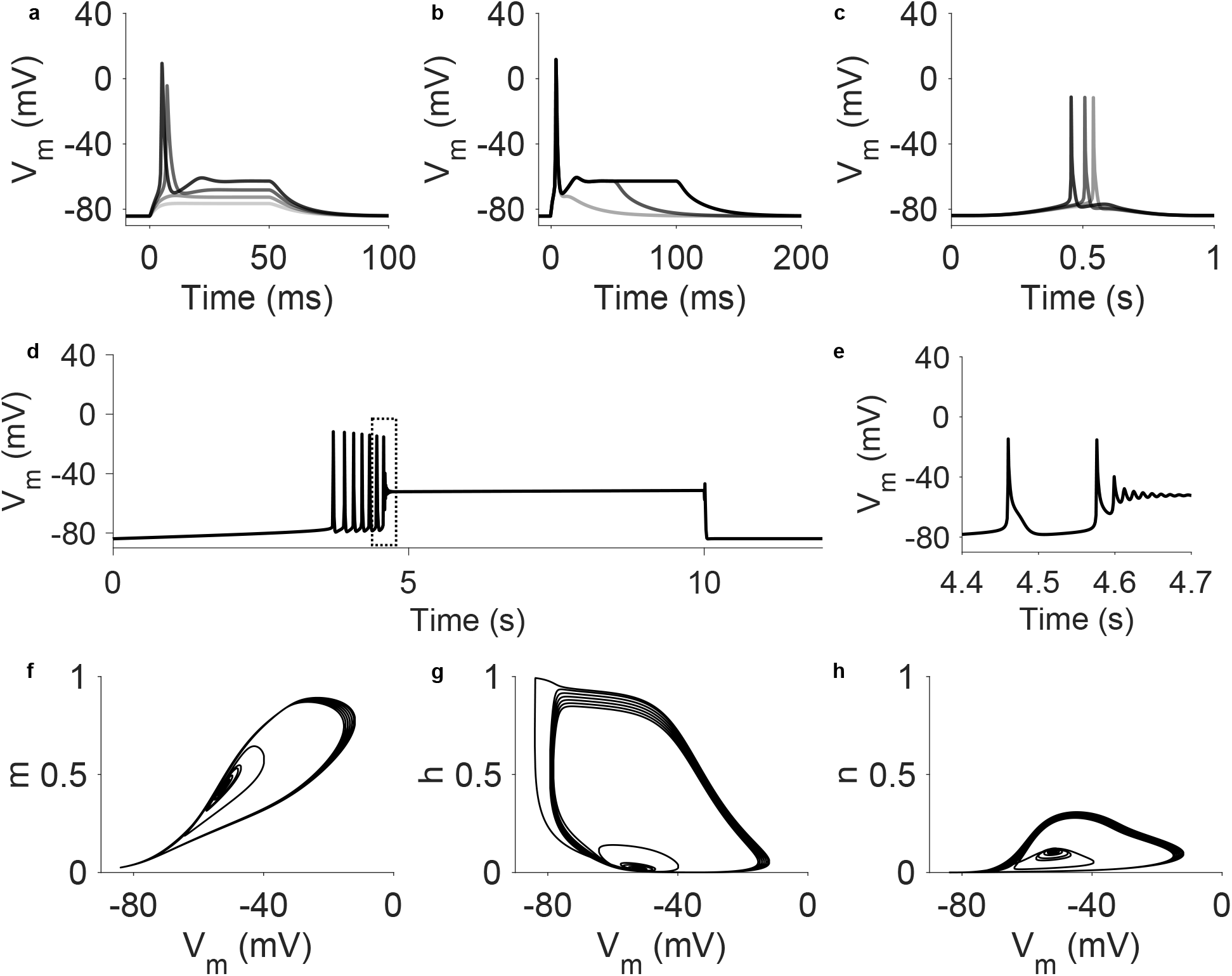
Hodgkin-Huxley model of muscle action potentials during optogenetic stimulation. **a, b** Simulated muscle action potentials with (**a**) 50 ms pulses with increasing ChR conductance (0.2 - 0.5 mS/cm^2^) and (**b**) pulses of constant ChR conductance (0.5 mS/cm^2^) with varying pulse durations 5, 50, and 100 ms. **c** Simulated muscle action potentials with 1 Hz sinuosoid stimulation of increasing ChR conductance (0.45 - 0.55 mS/cm^2^). **d** Simulated muscle action potentials with slowly ramping ChR conductance (0 - 0.125 mS/cm2) over 10 s with zoomed in view (**e**) at the transition between spiking and nonspiking behavior with vanishing amplitudes. **f-h** Hodgkin-Huxley gating parameters m, h, n as a function of V_m_ from simulation (**d**) shows disappearance of a stable limit cycle into a stable equilibrium, consistent with a supercritical Andronov-Hopf bifurcation.

## Notes

### Competing Interest Statement

The authors have declared no competing interest.

